# HPV Capsid-Derived Cationic Peptides for Cargo Delivery and Antiviral Activity

**DOI:** 10.64898/2026.05.06.723171

**Authors:** Vahagn Stepanyan, Silvia C. Finnemann, Patricio I. Meneses

**Author notes:** **Correspondence:** Patricio I. Meneses.

## Abstract

High-risk Human Papillomaviruses (HR-HPVs) are responsible for 5% of global cancers. While vaccines against HR-HPVs exist, there are no treatments available for individuals already infected. Cell-penetrating peptides (CPPs) have demonstrated antiviral properties against viruses by blocking viral entry and delivering antivirals into infected cells. Developing CPP-based therapies faces challenges including inefficient delivery of macromolecules and endosomal entrapment, which must be overcome for effective clinical application. This study identifies an HPV16 major capsid protein L1 derived cationic peptide as a potent CPP. Peptide uptake depended on both a cluster of cationic residues and the specific peptide sequence. Mechanistic studies showed peptide entry occurred via cell surface heparan sulfate-mediated, lipid-raft dependent endocytosis. The peptide efficiently delivered GFP into HaCaT keratinocytes, and associated with the Golgi apparatus, demonstrating endosomal escape. GFP fusion protein endocytosis relied on binding of the cationic peptide to cell surface heparan sulfates. Cell-penetrating ability was conserved among homologous regions of various HPV types. The peptide showed potent antiviral activity by inhibiting infection of HaCaT cells by several HR-HPV types collectively responsible for nearly all HPV-associated cancers. Excitingly, HPV18 L1-derived peptide from the homologous region exhibited potent antiviral activity against HPV16 by preventing viral internalization. Our findings characterize HPV-derived peptides as highly efficient CPPs with potential to deliver therapeutic agents into cells and assist in development of treatments for high-risk HPVs.

## 1 | Introduction

Human Papillomaviruses (HPVs) are nonenveloped double stranded DNA viruses that infect mucosal and cutaneous epithelium. HPVs measure ∼55 nm in diameter and are made up of 360 copies of the L1 major capsid proteins and up to 72 L2 minor capsid proteins. HPVs are divided into high-risk (HR) and low-risk (LR) types.^1,2^ HR-HPVs are responsible for approximately 5% of all global cancer cases. In 2020, there were an estimated 604,000 new cases of cervical cancer and 342,000 cervical cancer-related deaths worldwide. Nearly all cervical cancer cases are caused by HR-HPVs, with HPV16 and HPV18 accounting for 70% of these cases. The remaining cases are attributed to HPV31, 33, 45, 52, and 58.^3,4^ While effective preventive measures against high-risk HPVs-such as vaccinations-are available, there is currently no treatment for individuals already infected.^2,5,6^ Hence, there is a growing interest in peptide-based strategies, including the use of inhibitory, antimicrobial, and cell-penetrating peptides (CPPs) to block and treat existing viral infections.

CPPs are short, amphipathic or cationic peptides typically consisting of 6-30 amino acids.^7^ CPPs can be protein derived or synthetic, and they are characterized by their ability to cross the plasma membrane and deliver otherwise cell-impermeable molecules to cells. The first discovered and most widely studied CPP is TAT, derived from the HIV-1 TAT protein. The most widely studied synthetic CPPs are oligoarginines.^8–10^ CPPs have been used to successfully deliver a variety of macromolecules, including nucleic acids, siRNAs, proteins, and nanoparticles, into cells.^11–14^ Because of their ability to deliver bioactive compounds into cells, CPPs have become an attractive tool for delivering therapeutic agents to develop treatments for various conditions including cancer, neurodegenerative disease, inflammatory disease, cardiovascular disease and diabetes.^15–20^ While some CPPs have demonstrated antiviral properties, most lack intrinsic antiviral activity. Instead, their antiviral effect primarily stems from their ability to deliver antiviral compounds into cells, enabling the inhibition of a broad range of viral infections *in vitro* and *in vivo*. For example, CPP delivery of phosphorodiamidate morpholino oligomers (PMOs), antisense DNA oligonucleotides, have successfully inhibited HSV-1, Ebola virus, and Poliovirus in mice.^21–23^ Similarly, CPP-mediated siRNA delivery showed significant antiviral activity in vitro against HBV, HCV, and HIV-1 *in vitro* and *in vivo*.^24–26^

Most known CPPs are internalized via endocytosis. Although numerous CPPs have been developed and tested, many face significant challenges-particularly in achieving efficient intracellular delivery and facilitating endosomal escape, both of which are critical for the development of effective antiviral therapies. Various strategies have been deployed to overcome these limitations, such as dimerization and oligomerization of CPPs, which enhance delivery efficiency even at nanomolar concentrations and improve endosomal escape. For example, dimeric TAT peptide facilitated cargo escape from endosome.^27,28^ Another approach to improving CPP efficiency involved modulation of peptide stereochemistry. D-form oligoarginine peptide (D-R9) demonstrated higher cellular uptake than the L-form peptide (L-R9). Presence of both D- and L-Arginines resulted in higher delivery of siRNA into cells by chimeric 599 peptide.^29,30^ Cyclization of CPPs has shown to increase both cargo delivery and endosomal escape efficiency.^11,31^

Several methods have been employed to discover new and efficient CPPs, with the two primary approaches being trial-and -error and *in silico* design. The trial-and-error method typically involved experimentally testing peptides that share structural or physiochemical characteristics with well-known CPPs.^32^ In contrast, *in silico* design relies on machine learning algorithms trained on existing CPP datasets to predict novel candidates with high accuracy. However, a limitation of this approach is that non-CPP (negative) examples used for training are often random amino acid sequences. Some approaches utilize both *in silico* and trial-and-error to design new CPPs.^33–35^ For example, Wang et al. recently used MLCPP2.0, a machine-learning-based CPP prediction model, to estimate membrane penetration probability of peptides obtained from nuclear localization signal (NLS) database. Peptides with predicted probability uptake of greater than 50% were then tested for their cell-penetrating ability. As a result, the group found a CPP more efficient than TAT.^36^

We found previously that HPV16 L1 C-terminus-derived 15-amino-acid wild-type peptide (WT; 491-TSSTSTTAKRKKRKL-505), but not the scrambled peptide (SC; 491-SKTTKARLRTSKTKS-505), was able to inhibit HPV16 infection of HaCaT cells.^37^ Here, we report that the WT peptide acts as a potent CPP, whereas the SC peptide displays minimal activity. Through mutational analysis we found that peptide internalization is dependent on its basic residues and on the presence of cell surface heparan sulfates (HS). Functional analysis showed that the WT peptide internalized via lipid raft-mediated endocytosis. Moreover, the WT peptide could deliver GFP to the early endosome and showed localization with the Golgi apparatus. We also found that the CPP activity of the HPV16 L1 C-terminal peptide is conserved across multiple HPV types. Further analysis showed that the WT but not the SC peptide exhibited broad antiviral activity against the most common HR-HPV types. This antiviral property was also conserved in the homologous C-terminal region of the HPV18 L1 protein. Given the difference in internalization efficiency between the WT and SC peptides, we propose that both peptides can serve as useful templates for the rational design of next-generation CPPs. In particular, the WT peptide provides a model for efficient cellular uptake, while the SC peptide offers a comparative reference for identifying sequence features that limit internalization. Together, these peptides may guide the design of CPPs with improved delivery of bioactive compounds, such as antivirals, and enhanced endosomal escape. Furthermore, the broad intrinsic antiviral activity of the WT peptide highlights its potential as a candidate for antiviral development. Overall, these findings contribute to the ongoing development of peptide-based therapeutics for HR-HPV infections.

## 2 | Materials and Methods

### 2.1 | Cell culture

HaCaT cells, a spontaneously immortalized human epithelial cell line, AddexBio (San Diego, CA), HeLa cells (ATCC), and HEK-293TT cells (a kind gift from C. Buck, NIH, USA) were cultured in Dulbecco’s modified Eagle’s medium (DMEM; R&D Systems; Minneapolis, MN), supplemented with 10% heat inactivated fetal bovine serum (FBS) (GeminiBio; West Sacramento, CA).

### 2.2 | Peptide treatment

All peptides (Table 1) were obtained from LifeTein (USA). HaCaT or HeLa cells were seeded in 24-well plate at 75,000 cells/well. The next day the cells were treated with 4.5 μM indicated FITC-conjugated peptide for 2 h at 37 °C. Following the incubation the cells were either subjected to flow cytometry or confocal fluorescence microscope analysis.

**TABLE 1.**
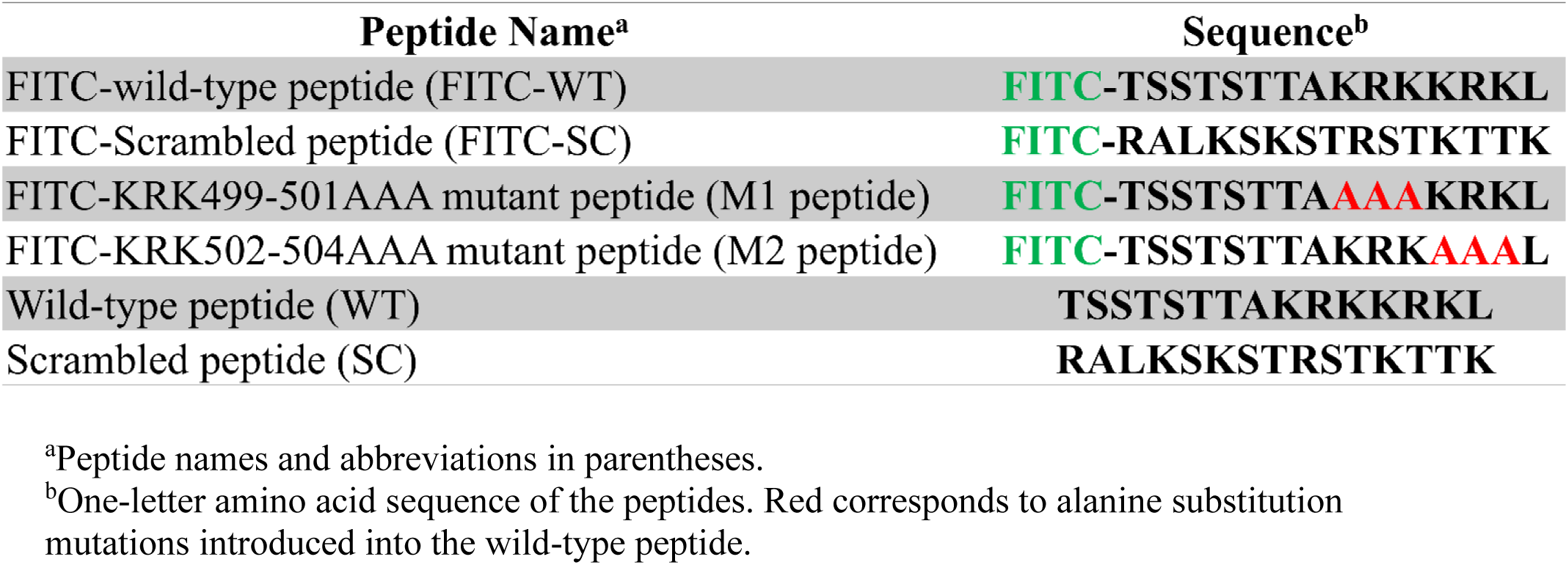
| Synthetic peptides used in this study

### 2.3 | GFP fusion protein construct, purification and treatment

pET His6 GFP TEV LIC cloning vector (1GFP) was a gift from Scott Gradia (Addgene plasmid # 29663 ; http://n2t.net/addgene:29663 ; RRID:Addgene_29663). Q5 site-directed mutagenesis kit (E0554S; New England BioLabs; Ipswich, MA) was used to insert the L1 C-terminal sequence into the plasmid. The PCR product was transformed into competent cells, and grown on Kanamycin agar plate, plasmid was purified from a colony that was grown from the plate. E. coli BL21(DE3) pLYsS cell (C2527H, New England BioLabs; Ipswich, MA) were transformed with new fusion GFP plasmids and grown on Kanamycin plates overnight at 37 °C. The following day a colony was picked and grown in Luria-Bertani (LB) broth overnight in a shaker at 250 rpm at 37 °C. The following day, once the OD=0.6, the expression of the plasmids was induced with isopropyl -β-D-thiogalactopyranoside (IPTG) at a final concentration of 1 mM for 2 h at 37 °C. The cells were spun down at 6000xg for 15 min. The bacteria cells were resuspended in 10mL of lysis buffer (50 mM NaH_2_PO_4,_ 300 mM NaCl, 10 mM Imidazole, Lysozyme 1 mg/mL) and incubated at room temperature for 30 min. The cells were then lysed using sonication, in presence of 1 mM Phenylmethylisulfonyl fluoride (PMSF). The lysates were centrifuged at 12,000xg for 20 min at 4 °C. Meanwhile, 500µL Ni-NTA agarose, for each construct, (30210, Qiagen; Germantown, MD) was resuspended in lysis buffer for 10 min and then spun down. The supernatant was transferred to new tubes with the 500µL Ni-NTA agarose and shaken for 30 min on ice. The agarose beads were spun down at 400xg for 3min, and resuspended in wash buffer (50 mM NaH_2_PO_4,_ 300 mM NaCl, 20 mM imidazole), transferred to chromatography column and rotated for 5 min at 4 °C. The column was washed two more times and finally eluted with elution buffer (50 mM NaH_2_PO_4,_ 300 mM NaCl, 300 mM imidazole). The purified proteins were dialyzed overnight in 1xPBS at 4 °C. The purified samples were then subjected to Coomassie blue staining and Western blot analysis.

HaCaT or HeLa cells were seeded in 24-well plate at 75,000 cells/well. The next day the cells were treated with 3 μM of GFP or GFP constructs for 2 h at 37 °C. Following the incubation the cells were either subjected to flow cytometry or confocal fluorescence microscope analysis. For endosomal escape, the cells were treated with 3 μM of GFP or GFP-16 and fixed after 12 h for confocal fluorescence microscopy.

### 2.4 | Endocytic inhibitor

HaCaT cells were seeded in 24-well plate at 75,000 cells/well. The next day the cell were treated with pre-treated with 20 μM chlorpromazine (Sigma Aldrich; St. Louis, MO, USA; C8313), 2 mM methyl-β-cyclodextrin (MβCD) (Sigma Aldrich; St. Louis, MO, USA; C4555), or 50 μM Dynasore (Sigma Aldrich; St. Louis, MO, USA; D7693) for 2 h, following the pre-incubation the cells were treated with 4.5 μM FITC-WT peptide, Human Transferrin CF488A (Biotium, 00081), or cholera toxin subunit B conjugated to Alexa Fluor-488 (Invitrogen, Thermo Fisher Scientific, C34775) in the presence of inhibitors for an additional 2 h. The cells were then trypsinized and subjected to flow cytometry.

### 2.5 | Heparin-agarose bead pulldown

GFP constructs were diluted to 50 ng/μL in 500 μL 1xPBS. Heparin agarose beads (Sigma Aldrich; St. Louis, MO, USA; H6508) were spun down and washed with 1xPBS, and 20 μL of agarose beads was added to each GFP construct and spun at room temperature for 1 h. The samples were spun down, washed and eluted with 500 mM NaCl for 5 min. Both, the input and the eluates were subjected to western blot analysis.

### 2.6 | Western Blot and Coomassie Blue staining

6X Laemmli buffer was added to purified GFP and eluted samples, boiled and subjected to SDS-PAGE. The resolved proteins were transferred to nitrocellulose membrane and blocked with 5% milk for 40 min. The membrane was washed with TBST (137mM NaCl, 2.7mM KCl, Tris-HCl pH 7.5, 0.1% Tween 20) and incubated with anti-6x His tag monoclonal antibody (Invitrogen, Thermo Fisher Scientific, MA1-21315). The membranes were visualized with LiCor Odyssey 9120 (LI-COR Biosciences; Lincoln, NE).

The purified samples were also subjected to Coomassie Blue staining. Purified GFPs were subjected to SDS-PAGE, followed by Coomassie Blue staining for 20 min at room temperature. The gels were then de-stained (50% methanol, 40% H_2_O, 10% acetic acid) and scanned with LiCor Odyssey 9120 (LI-COR Biosciences; Lincoln, NE).

### 2.7 | HPV pseudovirion (PsV) production and infection

HPV16, 18, 31, and 45 PsV were produced as described by Buck et al.. 293TT cells cultured in DMEM+10%FBS, were transfected with p16sheLL, p18sheLL, p31sheLL, and P45sheLL, a gift from John Schiller (Addgene plasmid # 37320 ; http://n2t.net/addgene:37320 ; RRID:Addgene_37320), (Addgene plasmid # 37321 ; http://n2t.net/addgene:37320 ; RRID:Addgene_37321), (Addgene plasmid # 37322 ; http://n2t.net/addgene:37322 ; RRID:Addgene_37322), (Addgene plasmid # 37323 ; http://n2t.net/addgene:37322 ; RRID:Addgene_37323), expressing HPV L1 and L2 capsid proteins, with reporter 8fwB plasmid, expressing GFP, serving as the pseudogenome. 48 h post transfection the 293TT cells were harvested, and PsVs were purified on OptiPrep gradient (27-39%) via ultracentrifugation at 50,000 rpm for 3.5 h at 16 °C using Beckman-Coulter L8-70M ultracentrifuge, and SW 55 Ti swinging-bucket rotor (Beckman Coulter; Brea, CA).

HaCaT cells were pre-chilled at 4 °C for 30 min., and 1500 viral genome equivalents (vge) per cell were added and allowed to bind for 2 h at 4 °C. Afterwards, excess PsVs was washed off three times with 1xPBS, followed by fresh DMEM+10% FBS addition. The cells were then treated with indicated concentration of peptides. After 48 h the cells were subjected to flow cytometry to determine the level of infection by detecting the number of GFP-positive cells.

### 2.8 | Flow cytometry

HaCaT cells were washed three times with 1xPBS, followed by trypsinization. The cells were spun at 600xg for 7 min, followed by three more washes. The cell pellets were then resuspended in 100 µL of 1xPBS followed by analysis with a BD Accuri C6 flow cytometer (BD Accuri C6; BD Biosciences; Franklin Lakes, NJ). 10,000 cells were analyzed per sample.

### 2.9 | Confocal laser scanning microscopy

HaCaT or HeLa cells were seeded in 24-well plate at 75,000 cells/well on glass coverslips. The following day they were subjected to either FITC-peptide or GFP construct treatment. At indicated time points the cells were fixed with 4% PFA for 5 min at room temperature. The cells were washed with 0.4% Trypan Blue to quench extracellular fluorescence of FITC and GFP. The cells were washed again three times with 1xPBS and permeabilized with 0.01% saponin for 15 min at room temperature. They were then stained with anti-EEA1 antibody EEA1 (C-15, sc-6414; Santa Cruz Biotechnologies) or anti-GM130 (P-20, sc-16268; Santa Cruz Biotechnologies) primary antibodies for 1 h, followed by 1xPBS wash. The cells were then incubated in anti-goat 647 secondary antibody for 30 min. followed by 1xPBS wash. The coverslips were mounted on slides with ProLong Gold Antifade reagent with DAPI (Invitrogen, Thermo Fisher Scientific, P36934). Primary antibodies were used at 1:100 dilution, and secondary antibodies at 1:2000 dilution. The slides were scanned using Leica TCS SP8 confocal microscope. ImageJ with JACop plugin was used for co-localization analysis.

## 3 | Results

### 3.1 | FITC-WT peptide internalization is dependent on the overall peptide sequence and the positively charged residues

We previously reported that WT but not the SC peptide inhibit HPV16 infection of HaCaT cells.^37^ Given this difference, we tested whether the WT or the SC peptide could function as a CPP. We obtained fluorescein isothiocyanate (FITC) conjugated WT and SC peptides, as shown in Table 1, and treated HaCaT cells with each peptide for 2 h, followed by flow cytometry to quantify internalization of each peptide. FITC-WT but not FITC-SC peptide efficiently entered HaCaT cells (Fig. 1A and B), and similar results were observed in HeLa cells (Fig. S1). Since cationic peptides, such as TAT, rely on a positively charged sequence for internalization we next determined if the WT peptide also relied on the cluster of positively charged sequences spanning 499-504 region.^38–40^ We obtained two FITC-conjugated M1 and M2 peptide, with 499-KRK-501 and 502-KRK-504 amino acids substituted with alanine residues (Table 1). HaCaT cells were treated with M1, M2 and FITC-WT peptides and subjected to flow cytometry. The results showed that the mutations in the positively charged residues significantly inhibited peptide internalization compared to the WT-FITC peptide (Fig. 1C and D).

**FIGURE 1.**
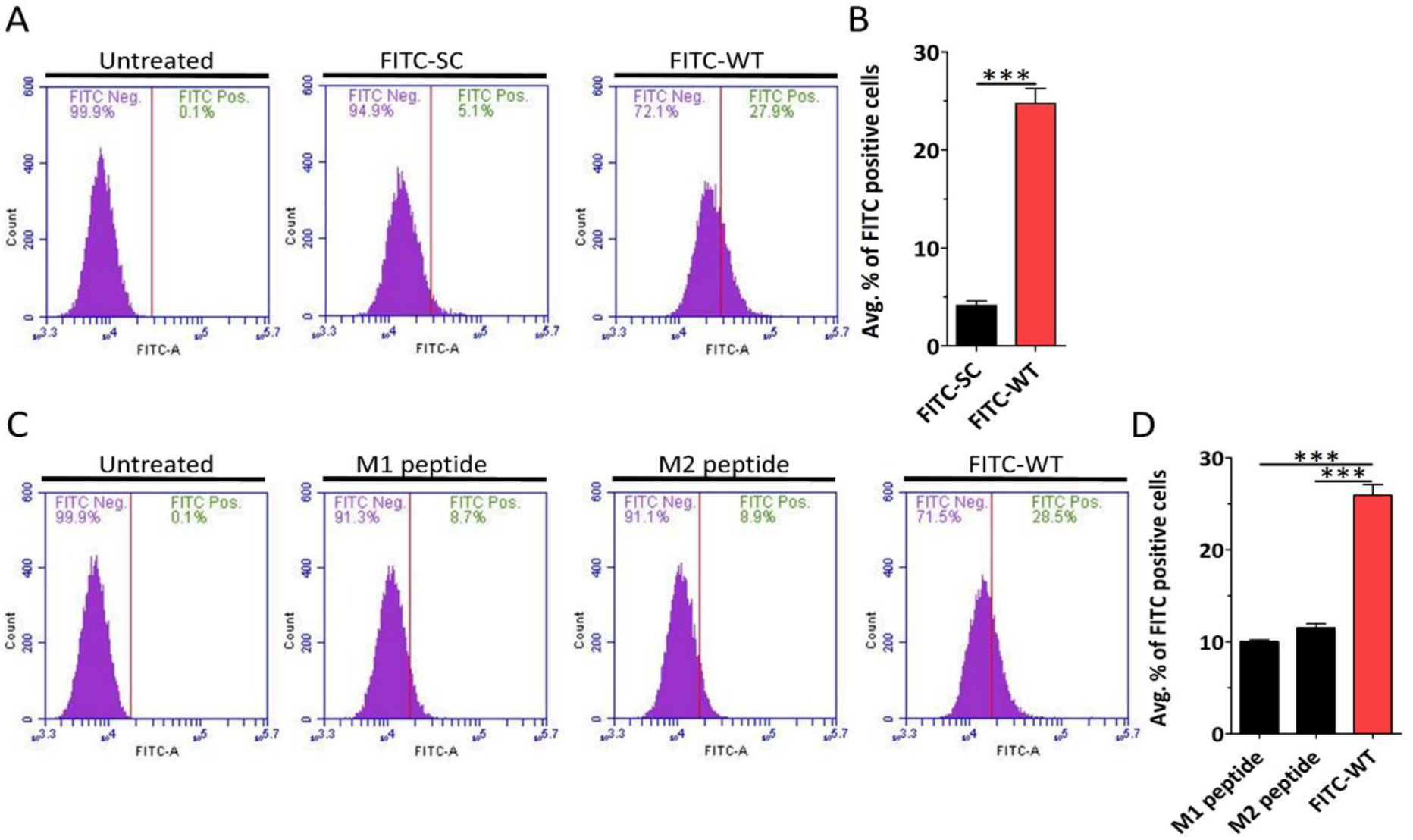
| WT peptide internalization is dependent on the overall peptide sequence and the positively charged residues. HaCaT cells were treated with 4.5 μM of the indicated peptide for 2 h and analyzed by flow cytometry. (A) Representative histograms show the percentage of FITC-positive HaCaT cells after treatment with FITC-SC and FITC-WT peptide. (B) Quantification of the average percentage. (C) Representative flow cytometry histogram showing the percentage of FITC-positive HaCaT cells after treatment with M1, M2, and FITC-WT peptides. (D) Quantification of the average percentage. Data are represented as mean ± SEM (n=3). Data were analyzed by unpaired two-tailed Student’s *t*-test; *, *P*<0.05; **, *P*<0.01; ***, *P*<0.001; *ns*, p>0.05.

### 3.2 | FITC-WT peptide enters cells via receptor-mediated lipid raft-dependent endocytosis

HPV16 and many cationic peptides rely on cell surface heparan sulfates for binding and subsequent entry into cells.^41,42^ As mutation of the positively charged sequences of the WT peptide inhibited peptide entry into cells, we set out to determine whether cell surface heparan sulfates (HS) mediated peptide entry.

HaCaT cells were treated with heparinase, fixed and stained for HS to confirm the removal from the cell surface (Fig. S3). After the removal of cell surface HS, HaCaT cells were treated with FITC-WT peptide. Flow cytometry analysis revealed removal of HS significantly decreased FITC-WT peptide internalization (Fig. 2A and B). These results revealed that WT-FITC peptide depended on HS receptors for internalization.

**FIGURE 2.**
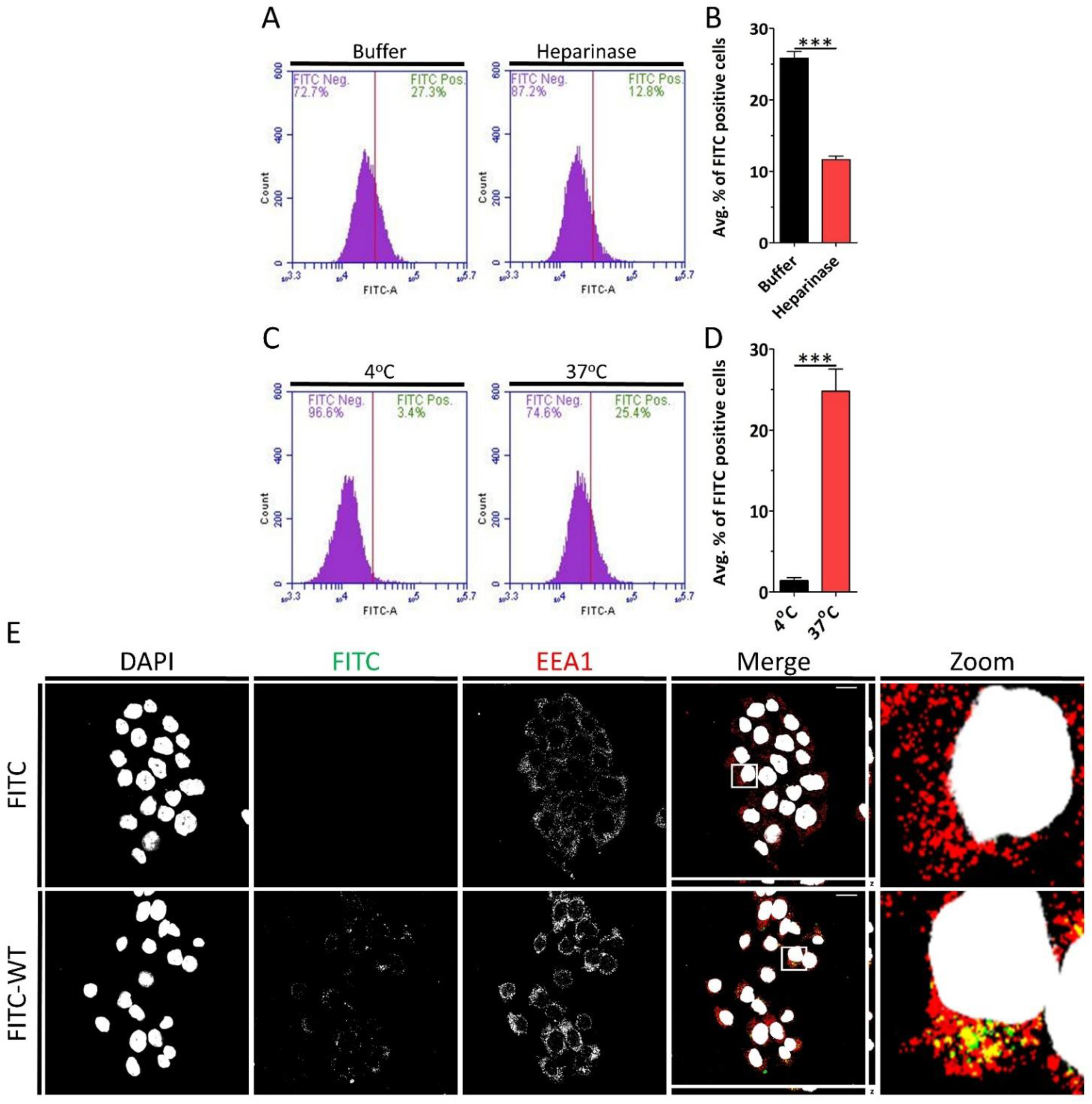
| FITC-WT peptide internalization is mediated by heparan sulfate via energy-dependent endocytosis. (A) HaCaT cells treated with Buffer (control) or Heparinase for 2 h at 37 °C. The cells were then treated with 4.5 μM FITC-WT for 2 h at 37 °C. Internalization was analyzed using flow cytometry. Representative histograms show the percentage of FITC-positive cells under each condition. (B) Quantification of the average percentage of FITC-positive HaCaT cells. (C) HaCaT cells were treated with 4.5 μM FITC-WT for 2 h at 4 °C or 37 °C, followed by analysis via flow cytometry. Representative histograms show the percentage of FITC-positive cells. (D) Quantification of the average percentage of FITC-positive HaCaT cells at 4 °C and 37 °C after 2 h. (E) Confocal fluorescence microscopy images of HaCaT cells treated with 4.5 μM FITC or FITC-WT for 2 h at 37 °C. Gray corresponds to DAPI (nuclear stain), green corresponds to FITC-WT, and red corresponds to EEA1 (early endosomal marker). Scale bar: 20 μm. Data are represented as mean ± SEM (n=3). Data were analyzed by unpaired two-tailed Student’s *t*-test; *, *P*<0.05; **, *P*<0.01; ***, *P*<0.001; *ns*, p>0.05.

CPPs may utilize various mechanisms for cell entry. Some peptides internalize via energy-independent pathways and others, more commonly, through energy-dependent endocytosis.^42,43^ To determine the internalization mechanism of FITC-WT peptide into HaCaT cells, we first chilled cells at 4 °C followed by addition of FITC-WT peptide. The cells were then either left at 4°C or transferred to 37°C. 2 hours later, cells were subjected to flow cytometry, which revealed that the peptide internalized in an energy dependent manner (Fig. 2C and D), possibly via endocytosis. To determine whether FITC-WT peptide internalization occurred via endocytosis, we treated HaCaT cells with FITC or FITC-WT peptide for 2 h, following which the cells were fixed and stained for early endosomal antigen 1 (EEA1), an early endosomal marker. Confocal microscopy showed co-localization of FITC-WT peptide with EEA1 (Fig. 2E) indicating that peptide internalization occurred via energy-dependent endocytosis.

We next used standard pharmacological endocytic inhibitors to identify the specific endocytic pathway utilized by the peptide. Because clathrin-mediated endocytosis (CME) is the most extensively studied endocytic pathway, we decided to determine its role in the peptide internalization.^44^ We pre-treated HaCaT cells with chlorpromazine, a widely used CME inhibitor, for 2 h followed by addition of the WT-FITC peptide for an additional 2 h in the presence of the inhibitor. Alexa Fluor 488-conjugated transferrin was used as positive control for CME.^45^ Flow cytometry analysis showed that chlorpromazine significantly increased FITC-WT peptide internalization. As anticipated, we observed a decrease in transferrin internalization (Fig 3A and B). These results indicated that the peptide internalized via CME-independent pathway. Next, we tested whether caveolae-dependent endocytosis mediated peptide internalization. We pre-treated HaCaT cells with methyl-β-cyclodextrin (MβCD), a widely used caveolae-mediated endocytosis inhibitor. Following pre-incubation, HaCaT cells were treated with either FITC-WT peptide or Alexa Fluor 488-conjugated cholera toxin subunit B (CTB), a positive control for caveolae-mediated endocytosis.^46,47^ Flow cytometry analysis showed that MβCD reduced FITC-WT peptide internalization. As anticipated, we observed a decrease in internalization of CTB (Fig. 3C and D). As MβCD functions by depleting cholesterol from cell membrane, it has an inhibitory effect on both caveolae-mediated endocytosis as well as lipid raft-dependent endocytosis.^46,48^ However, only caveolae-dependent endocytosis is also dynamin-dependent. We thus used the dynamin inhibitor dynasore to determine which of the two pathways was utilized by the peptide.^49^ We pre-treated cells with dynasore, followed by treatment with FITC-WT peptide or transferrin as positive control. Flow cytometry analysis showed a significant increase in FITC-WT internalization, whereas the expected decrease in transferrin was observed (Fig. 3E and F). Taken together, these results indicated that the WT-FITC peptide internalized in clathrin-, caveolae-, and dynamin-independent but lipid raft-dependent endocytosis. Notably, none of the inhibitors showed cytotoxic effects at the concentration used in the experiments (Fig. S4A).

**FIGURE 3.**
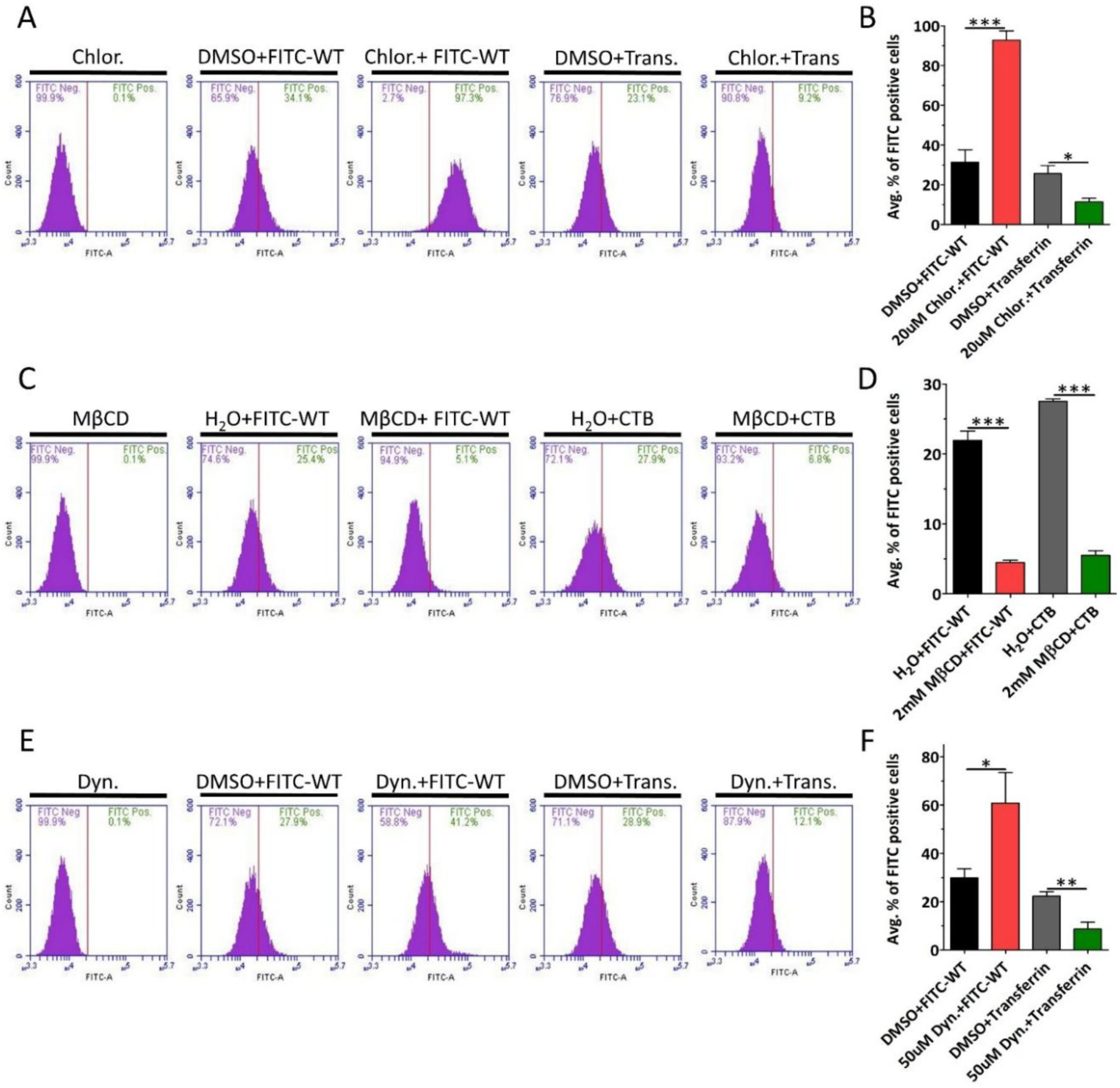
| FITC-WT peptide internalizes via lipid-raft endocytosis, independent of Clathrin, Caveolae, and Dynamin. (A) HaCaT cells were either pretreated with vehicle, DMSO, or 20 μM chlorpromazine, followed by treatment with either FITC-WT peptide or transferrin, positive control, for 2 h at 37 °C in the presence of the inhibitor. Internalization was analyzed using flow cytometry. Representative histograms show the percentage of FITC-positive cells. (B) Quantification of the average percentage of FITC-positive cells. (C) HaCaT cells were either pretreated with vehicle, H_2_O, or 2 mM MβCD, followed by treatment with either FITC-WT peptide or CTB, positive control, for 2 h at 37 °C in the presence of the inhibitor. Internalization was analyzed via flow cytometry. Representative histograms show the percentage of FITC-positive cells. (D) Quantification of the average percentage of FITC-positive cells. (E) HaCaT cells were either pretreated with vehicle, DMSO, or 50 μM dynasore, followed by treatment with either FITC-WT peptide or transferrin, positive control, for 2 h at 37 °C in the presence of the inhibitor. Internalization was analyzed via flow cytometry. Representative histograms show the percentage of FITC-positive cells. (F) Quantification of the average percentage of FITC-positive cells. Data are represented as mean ± SEM (n=3). Data were analyzed by unpaired two-tailed Student’s *t*-test; *, *P*<0.05; **, *P*<0.01; ***, *P*<0.001; *ns*, p>0.05.

### 3.3 | GFP-WT peptide enters HaCaT cells

One of the defining characteristics of CPPs is their ability to deliver cargo into cells that, on their own, cannot enter cells. These include macromolecules such as proteins, DNA and RNA as well as therapeutic agents such as antiviral and antibacterial drugs.^11,12,22,50^ Here, we created and purified GFP and WT peptide (GFP-WT) fusion proteins (Fig. 4A). HaCaT cells were treated with either GFP or GFP-WT proteins. The cells were then subjected to flow cytometry and revealed a modest but significant increase in GFP-WT internalization (Fig 4B and C). We suspected that GFP might be interfering with the uptake capacity of the peptide. Cargoes have been shown to decrease the efficiency of CPPs.^51–56^ To address this, we increased the peptide length to include the last 30 amino acids of the L1 protein, spanning residues 478-505. We created and purified this new GFP-peptide construct, referred to as GFP-16 (Fig. 4D), and treated HaCaT cells with either GFP or GFP-16. Flow cytometry analysis showed a marked increase in GFP-16 internalization into HaCaT cells (Fig. 4E and F). Confocal microscopy analysis showed that GFP-16 co-localized with EEA1 (Fig. 4G), suggesting that GFP-16 internalized via endocytosis. Similar results were observed in HeLa cells (Fig. S2). A major shortcoming of CPP cargo delivery is endocytic entrapment of the CPP.^57^ We incubated HaCaT cells with GFP or GFP-16 for 12 h, allowing sufficient time for potential escape. Confocal microscopy analysis revealed GFP-16 co-localized with the *cis*-Golgi apparatus marker GM130 indicating GFP-16 endosomal escape (Fig. 4H).

**FIGURE 4.**
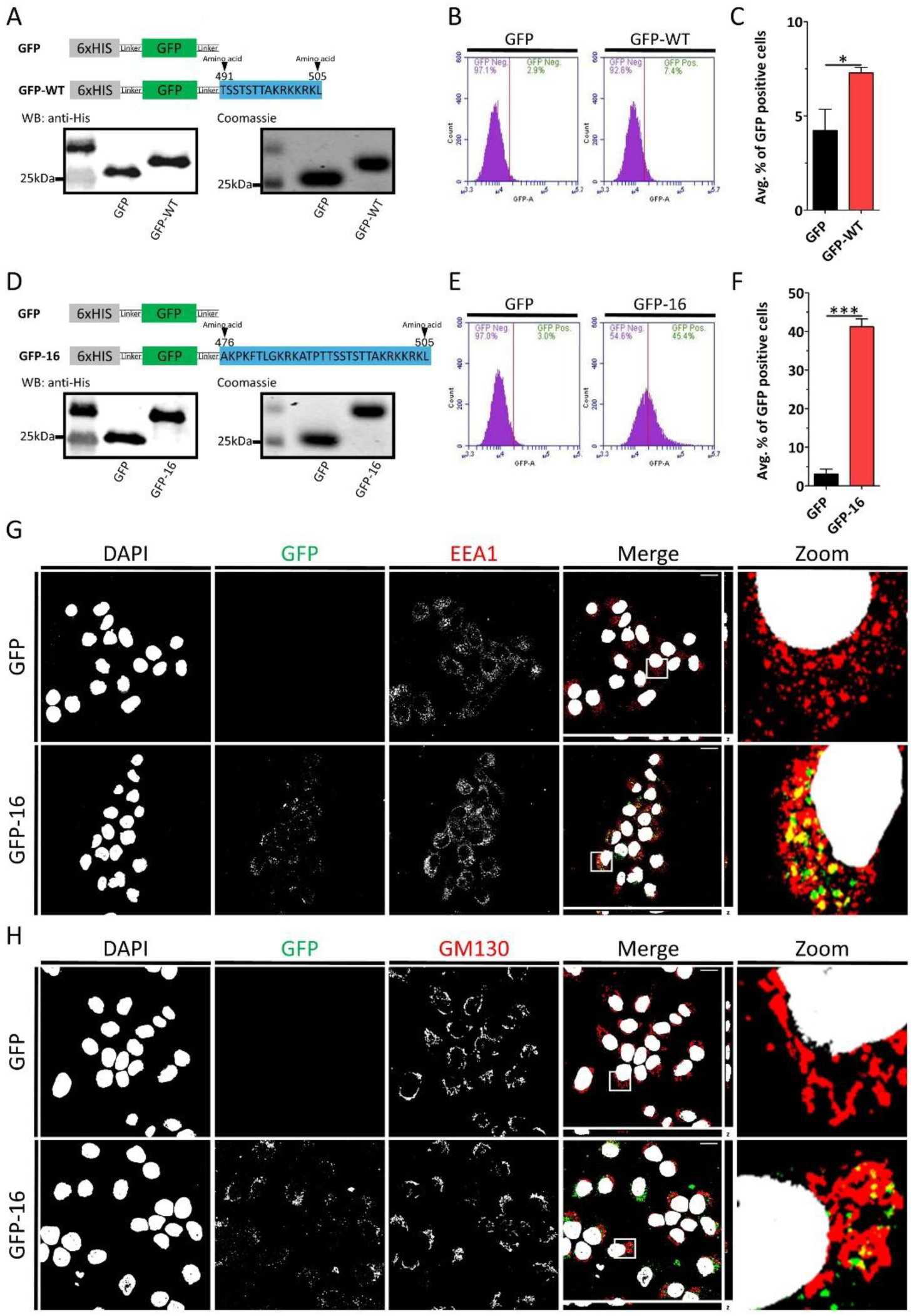
| GFP-WT peptide enters HaCaT cells. (A) Schematic representation of GFP and GFP-WT fusion protein construct. Below are the Western blot and Coomassie blue-stained SDS-PAGE gel of purified GFP. (B) HaCaT cells were treated with 3 μM of GFP or GFP-WT at 37 °C for 2 h. Internalization was analyzed using flow cytometry. Representative histograms show the percentage of GFP-positive cells. (C) Quantification of the average percentage of GFP-positive cells. (D) Schematic representation of GFP and GFP-16 fusion protein construct. Below are the Western blot and Coomassie blue-stained SDS-PAGE gel of purified GFP. (E) HaCaT cells were treated with 3 μM of GFP or GFP-16 at 37 °C for 2 h. Internalization was analyzed using flow cytometry. Representative histograms show the percentage of GFP-positive cells. (F) Quantification of the average percentage of GFP-positive cells. (G) Confocal fluorescence microscopy images of HaCaT cells treated with 3 μM GFP or GFP-16 for 2 h at 37 °C. Gray corresponds to DAPI (nuclear stain), green corresponds to GFP, and red corresponds to EEA1 (early endosomal marker). (H) Confocal fluorescence microscopy images of HaCaT cells treated with 3 μM GFP or GFP-16 for 12 h at 37 °C. Gray corresponds to DAPI (nuclear stain), green corresponds to GFP, and red corresponds to GM130 (*cis*-Golgi marker). Scale bar: 20 μm. Data are represented as mean ± SEM (n=3). Data were analyzed by unpaired two-tailed Student’s *t*-test; *, *P*<0.05; **, *P*<0.01; ***, *P*<0.001; *ns*, p>0.05.

### 3.4 | GFP-16 endocytosis depends on interaction of 499-KRKKRK-505 residues with cell surface heparan sulfate

To determine whether GFP-16 relied on HS like WT peptide, we removed cell surface HS by treating HaCaT cells with heparinase (Fig. S5). After HS removal, the cells were treated with GFP-16. Flow cytometry analysis showed that GFP-16 internalization was strongly dependent on cell surface HS (Fig. 5A and B). As we observed previously, the FITC-WT peptide internalization also depended on positively charged residues in the 499-505 region. To determine whether GFP-16 also relied on this cluster of positively charged residues, we substituted them with alanine via site-directed mutagenesis, referred to as KRKKRK499-505AAAAAA. We then created and purified this mutant GFP along with GFP-16 (Fig. 5C). HaCaT cells were treated with GFP-16 or KRKKRK499-505AAAAAA mutant and analyzed by flow cytometry. KRKKRK499-505AAAAAA mutation significantly reduced GFP internalization in HaCaT cells compared to GFP-16 (Fig. 5D and E). Given that we observed that GFP-16 internalization depends on both cell surface HS and the positively charged amino acid residues, we sought to determine whether these residues directly interact with HS. To test this, we performed heparin-agarose pull-down assays with GFP-16 and the KRKKRK499-505AAAAAA mutant confirming direct heparin binding of GFP-16 but not the mutant GFP (Fig. 5F).

**FIGURE 5.**
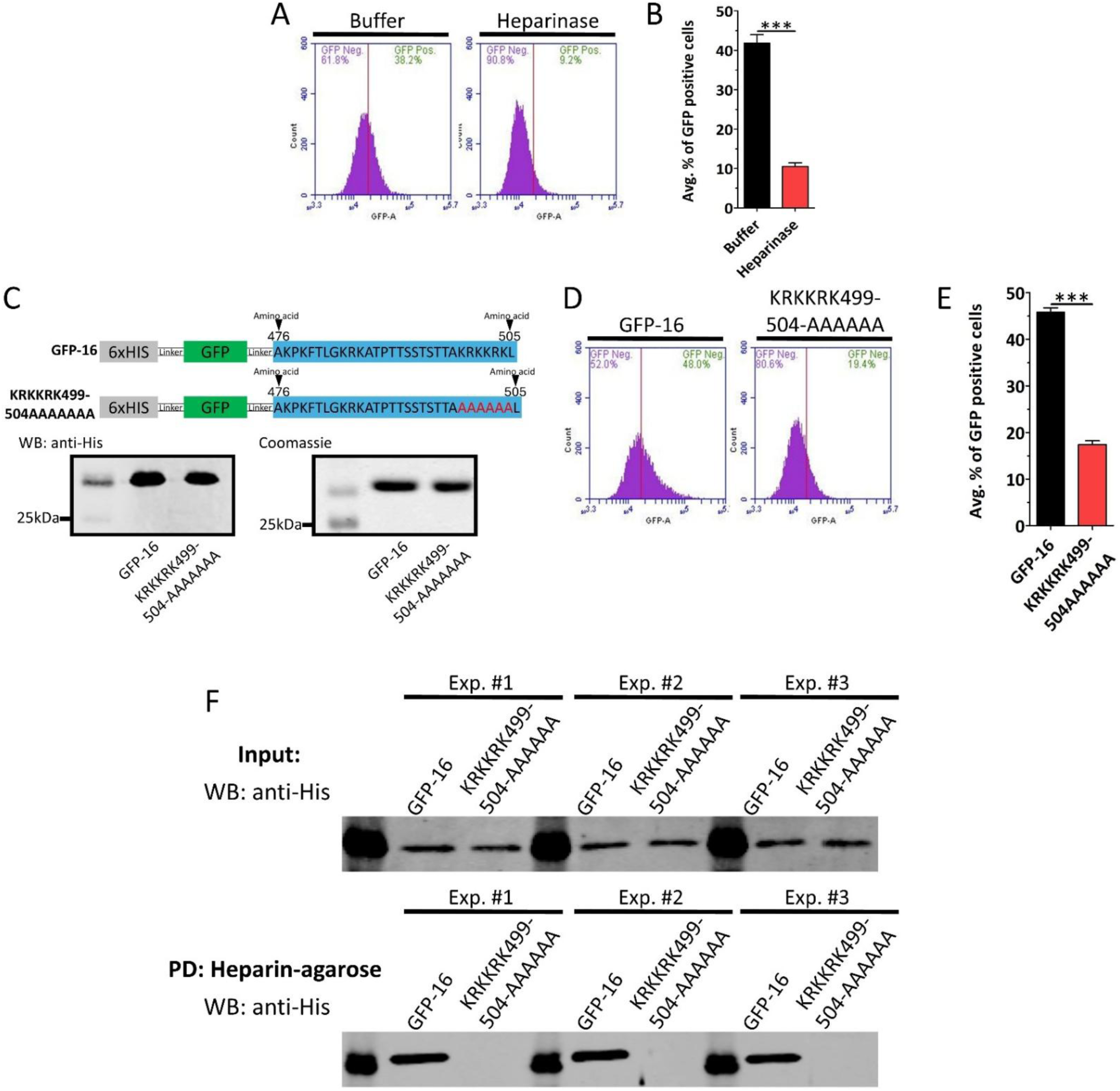
| GFP-16 endocytosis depends on interaction of 499-KRKKRK-505 residues with cell surface heparan sulfate. (A) HaCaT cells treated with Buffer (control) or Heparinase for 2 h at 37 °C. After the treatment, the cells were treated with 3 μM GFP-16 for 2 h at 37 °C. Internalization was analyzed using flow cytometry. Representative histograms show the percentage of GFP-positive cells under each condition. (B) Quantification of the average percentage of GFP-positive cells. (C) Schematic representations of GFP-16 and GFP-KRKKRK499-504AAAAA mutant fusion protein constructs. Below are the Western blot and Coomassie blue-stained SDS-PAGE gel of purified GFP. (D) HaCaT cells were treated with 3 μM of GFP-16 and GFP-KRKKRK499-504AAAAA at 37 °C for 2 h. Internalization was analyzed using flow cytometry. Representative histograms show the percentage of GFP-positive cells. (E) Quantification of average percentage of GFP-positive cells. (F) GFP-16 and KRKKRK499-504AAAAA mutant were incubated with heparin-agarose beads. The samples were spun down, washed with 1xPBS, and eluted with 500 mM NaCl. Western blot of the input sample (top). Western blot of the eluate sample (bottom). Scale bar: 20 μm. Data are represented as mean ± SEM (n=3). Data were analyzed by unpaired two-tailed Student’s *t*-test; *, *P*<0.05; **, *P*<0.01; ***, *P*<0.001; *ns*, p>0.05.

### 3.5 | Cell-penetrating activity of the L1 C-terminus is conserved across high- and low-risk HPV types

All HPVs have a positively charged L1 C-terminus that serves as nuclear localization signal. However, the C-terminal region is amongst the least conserved regions across HPV types.^58^ We sought to determine whether these sequences might be functionally conserved, specifically whether they can act as CPPs, like the C-terminal sequence of HPV16 L1. To this end, we selected C-terminal regions from three high-risk HPV types-45, 31, and 18- which, together with HPV16, account for most of the HPV-related cancers, as well as from low-risk type 6.^4,6^ We generated and purified these GFP peptide fusion constructs, referred to as GFP-45, GFP-31, GFP-18, and GFP-6 (Fig. 6A). HaCaT cells were treated with GFP, GFP-45, -31, -18 or -6. Flow cytometry analysis revealed that all peptides successfully delivered GFPs into HaCaT cells. Notably, GFP-6 exhibited the highest CPP activity, delivering GFP into more than 90% of cells. (Fig. 6B and C). Confocal microscopy showed that all GFP constructs colocalized with EEA1 (Fig. 6D), suggesting that they internalized via endocytosis.

**FIGURE 6.**
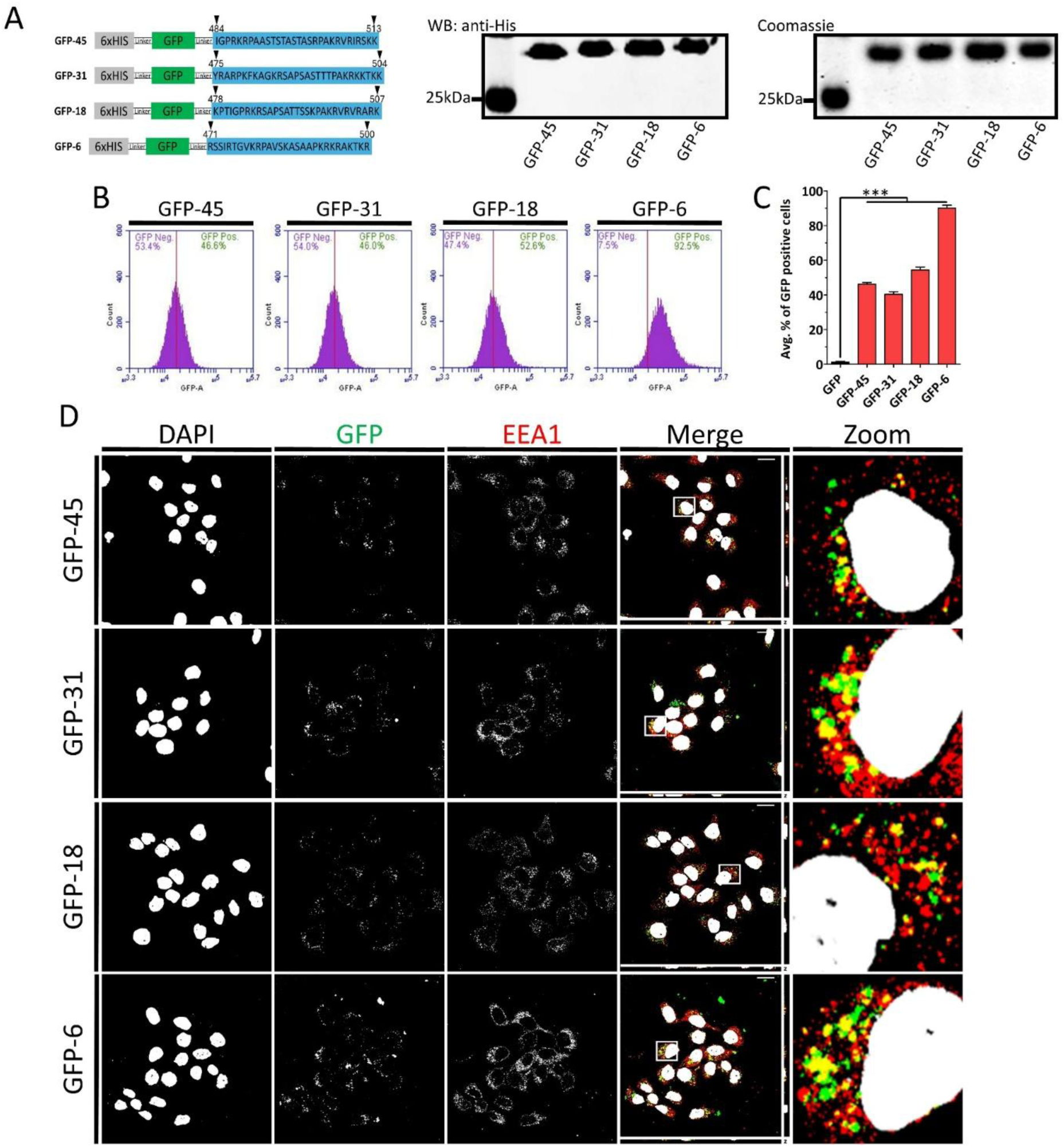
| Cell-penetrating activity of the L1 C-terminus is conserved across high- and low-risk HPV types. (A) Schematic representation of GFP-45, -31, -18 and -6 fusion protein constructs. To the right are the Western blot and Coomassie blue-stained SDS-PAGE gel of purified GFPs. (B) HaCaT cells were treated with 3 μM of GFP-45, -31, -18 or -6 at 37 °C for 2 h. Internalization was analyzed using flow cytometry. Representative histograms show the percentage of GFP-positive cells. (C) Quantification of the average percentage of GFP-positive cells. (D) Confocal fluorescence microscopy images of HaCaT cells treated with 3 μM GFP or GFP-16 for 2 h at 37 °C. Gray corresponds to DAPI (nuclear stain), green corresponds to GFP, and red corresponds to EEA1 (early endosomal marker). Scale bar: 20 μm. Data are represented as mean ± SEM (n=3). Data were analyzed by one-way ANOVA (Bonferroni’s multiple comparisons test; *, P<0.05; **, P<0.01; ***, P<0.001; ns, p>0.05).

### 3.6 | HPV16 C-terminal WT peptide acts as a potent antiviral

The HPV16-derived WT peptide spanning 491-505 region inhibits HPV16 infection.^37^ Here, we investigated whether the WT peptide would have the same effect on HPV18, 31, and 45 infection. HaCaT cells were incubated with HPV16, 18, 31 or 45 to allow virus to bind. Excess virus was then washed off, and fresh media was added along with either SC or WT peptide for 48 h followed by flow cytometry analysis. The WT peptide significantly inhibited infection of HaCaT cells by HPV16, 18, 31, and 45, while the SC peptide had no effect (Fig. 7A).

**FIGURE 7.**
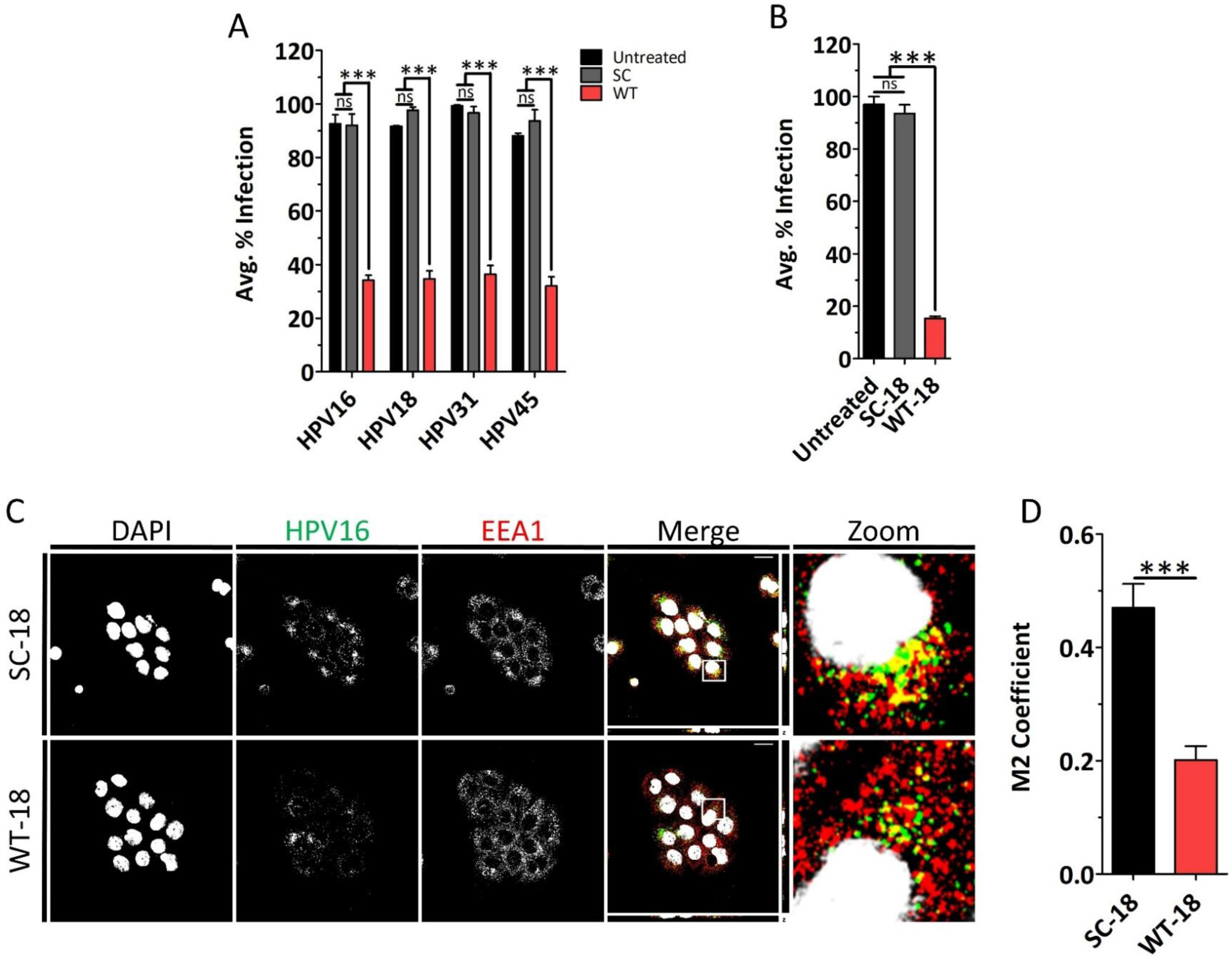
| HPV16 C-terminal WT peptide acts as a potent antiviral. (A) The WT but not the SC peptide inhibits infection of HaCaT cells by the most common high-risk HPV types. HaCaT cells were incubated with HPV 16, 18, 31, or 45 at 4 °C for 2 h. Excess virus was washed off by 1X PBS, and fresh media was added. Cells were treated with 18 μM SC or WT peptide. After 48 h, the infection level was assessed using flow cytometry. For each, results were normalized to the sample with the highest infection. (B) HaCaT cells were incubated with HPV16 4 °C for 2 h. Excess virus was washed off by 1X PBS, and fresh media was added. Cells were treated with 18 μM SC-18 or WT-18 peptide. After 48 h, the infection level was assessed by flow cytometry. Results were normalized to the sample with the highest infection. (C) Confocal fluorescence microscopy images of HaCaT cells showing co-localization of HPV16 with early endosome at 2 h post 18 μM SC-18 or WT-18 peptide treatment. (D) Quantification of early endosomal co-localization with HPV16. Scale bars indicated are 20 μm. Data are represented as mean ± SEM (n=3). Data were analyzed by one-way ANOVA (Bonferroni’s multiple comparisons test) and by unpaired two-tailed Student’s *t*-test; *, P<0.05; **, P<0.01; ***, P<0.001; ns, p>0.05.

As HPV18 is the second most common HR-HPV type, we also obtained a WT peptide from the HPV18 homologous region spanning amino acids 493-507 of the L1 protein, referred to as WT-18. We also obtained SC peptide composed of the same amino acids but in a random order, referred to as SC-18 (Table 1). We then tested WT-18 and SC-18 inhibitory activity against HPV16, as it is the most common HR-HPV type. Flow cytometry analysis showed that WT-18 had a potent inhibitory effect on HPV16 infection, while SC-18 had no effect (Fig. 7B). Confocal microscopy co-localization analysis further confirmed diminished EEA1 co-localization with HPV16 (Fig. 7C and D). None of the peptides had cytotoxic effects (Fig. S4B). Altogether, we found that the WT peptide derived from HPV16 had broad antiviral activity against several HR-HPV types and the WT-18 peptide derived from HPV18 inhibited HPV16 infection. These results suggest that the function of the L1 C-termini is conserved across HR-HPV types.

## 4 | Discussion

Plasma membranes present a significant barrier to the intracellular delivery of hydrophilic macromolecules.^59^ CPPs can overcome this limitation by crossing the plasma membrane and, more importantly, delivering biologically active cargo such as proteins, siRNA, DNA, and drugs into cells. Among CPPs, cationic peptides are the most well-studied class, including well-known examples like TAT and octaarginine peptides. These peptides are rich in arginine residues, which are believed to play a central role in their membrane translocating ability.^8–10^ Substituting arginine residues has been shown to reduce peptide efficiency. Because of the importance of arginine residues, CPPs are sometimes referred to as arginine-rich peptides. ^38–40^ When designing new CPPs, candidate sequences are frequently based on arginine-rich motifs, often including NLS. New *in silico* methods, which are based on machine learning, are trained on well-known CPPs, and on this basis they are taught to identify new CPPs. ^32,34–36^ We demonstrated that a peptide derived from the NLS of the HPV16 L1 major capsid protein functions as a CPP, with its activity dependent on positively charged 499-KRK-501 and 502-KRK-505 residues (Fig. 1C and D). Moreover, we found that the scrambled peptide, which retains the overall positive charge, showed poor internalization. These results suggest that the specific amino acid sequence is also critically important for cationic peptide internalization. We propose that both WT and SC peptides can serve as valuable templates for *in silico* CPP design. Instead of relying solely on comparisons between known CPPs and non-CPPs, these models could be trained using highly efficient CPPs, such as the WT peptide, and less efficient variants, such as the SC peptide. This approach has the potential to improve the identification and rational design of novel, more efficient CPPs.

We found that removal of cell surface HS significantly reduced internalization of WT peptide compared to buffer-treated control cells.^41,42^ Given the poor internalization of the SC peptide, it is likely that the interaction between the WT peptide and HS is sequence-specific. Future studies should explore potential endocytic receptors, such as syndecans (SDCs). Letoha et al. showed that both TAT and R_8_ peptides prefer SDC4 over SDC1 and SDC2 for binding and internalization. ^41^ Since a limitation of CPPs is their lack of tissue specificity, identifying a preferred receptor for HPV WT peptide may enable tissue-specific targeting of the peptide.

Our experiments also reveal that the WT peptide entry pathway is lipid raft-dependent, but clathrin-, caveolae-, and dynamin-independent endocytosis. While these results establish the L1 C-terminal-derived peptide as a CPP, we sought to determine whether it can perform one of the central functions of CPPs, namely the delivery of cargo into cells. To test this, we created GFP fusion proteins. However, because cargo is known to affect CPP efficiency, extension of the peptide length to GFP-16 including residues 476-505 of the L1 C-terminus was necessary to yield efficient delivery into HaCaT cells.^7,52–56^ Endocytosed GFP-16 eventually reached the Golgi apparatus demonstrating endosomal escape. Peptides capable of escaping the endosome can be used to deliver therapeutic agents, such as drugs and siRNA, into cells.^60^ Thus, the HPV-derived WT peptide may be capable of delivering these therapeutic agents into cells. Our results provide a foundation for the development of L1 C-terminus-based delivery peptides in therapeutic applications. Future studies are needed to elucidate the molecular mechanism underlying its endosomal escape and trafficking.

Mutation of the positively charged residues of GFP-16, KRKKRK499-505AAAAAA, demonstrated that this region plays a primary role in GFP internalization through direct binding of these residues to cell surface HS. Despite the increased length of the peptide, the cluster of positively charged residues still played a primary role in GFP fusion protein internalization, similar to the WT peptide. This finding further supports the notion that the peptide’s interaction with HS is specific and may involve obligate, yet unidentified, receptor.

Interestingly, HPV L1 C-terminus region is among the least conserved in the L1 protein.^61^ However, we found that all five HPV L1 C-terminal peptides we tested acted as CPP when fused to GFP, with the HPV6 C-terminal peptide showing particularly strong CPP activity. These results suggest strong functional conservation of the C-terminal region across diverse HPV types and indicate that HPV L1 C-termini represent a rich source of CPPs. Although we saw an endosomal escape with HPV16 L1 C-terminal peptide, similar assays are necessary for GFP-45, -31, -18, and -6 peptides. A recent study showed that cyclization of CPPs such as TAT and R_8_ peptides increased their efficiency by several fold. ^11^ Thus, similar peptide modifications may enhance both cellular entry and endosomal escape of the peptides in this study.

Finally, we found that the WT peptide, but not the SC peptide, exhibits broad inhibitory activity to all HPV types tested. Similarly, the HPV18 derived peptide inhibits HPV16 infection by preventing internalization as confocal microscopy results showed. These results indicate that HPV L1 C-terminus-derived peptides possess antiviral activity across HPV types. While CPPs have utility for delivery of antiviral cargos into cells, our data suggest these peptides themselves are biologically active both as CPPs and as antiviral agents. ^21–23^ As neither SC nor SC-18 peptides showed any antiviral activity, this supports a mechanism involving sequence specific binding. We propose that WT and WT-18 peptides inhibit infection via competitive binding to a receptor, thereby preventing viral internalization.

## 5 | Conclusion

Our study demonstrates that HPV L1 C-terminus-derived peptides possess significant cell-penetrating and antiviral activities, mediated through specific interaction with cell surface HS and efficient endosomal escape. These findings not only expand the repertoire of functional CPPs but also highlight the therapeutic potential of these peptides as delivery vehicles and antiviral agents. Future research should focus on optimizing their specificity and efficacy for clinical applications. Together, these efforts could contribute to the development of novel strategies for targeted therapeutic delivery and HR-HPV infection treatment.

## Acknowledgments

We would like to thank Dr. Buck and Dr. Christensen for providing essential reagents.

