## Supplementary Figures S1-S5 for "HPV Capsid-Derived Cationic Peptides for Cargo Delivery and Antiviral Activity"

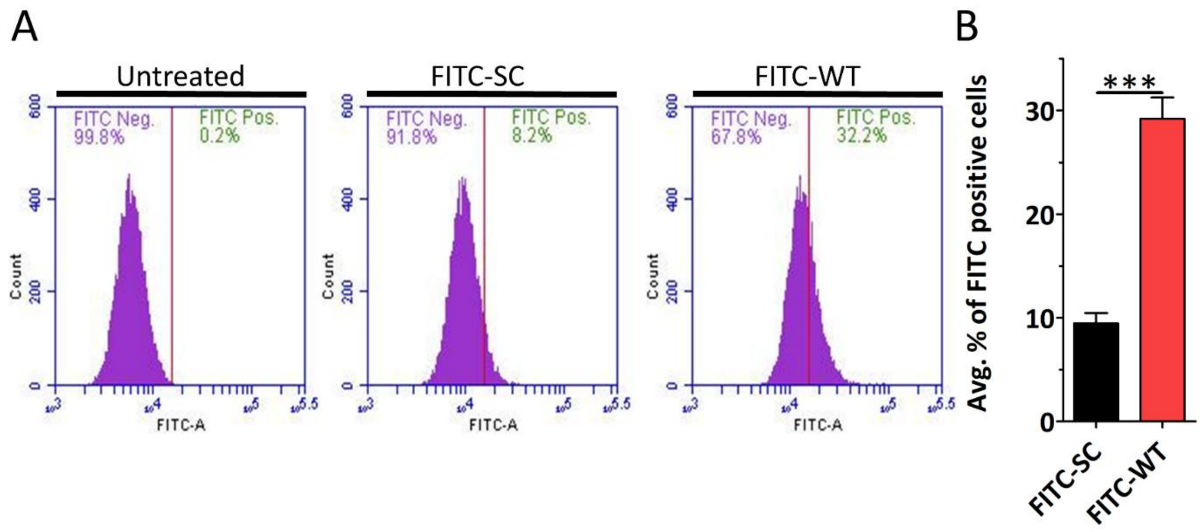


**FIGURE S1 |** WT peptide internalization is dependent on the overall peptide sequence. HeLa cells were treated with 4.5 µM of the indicated peptide for 2 h and analyzed by flow cytometry. (A) Representative histograms show the percentage of FITC-positive HeLa cells after treatment with FITC-SC and FITC-WT peptide. (B) Quantification of the average percentage. Data are represented as mean ± SEM (n=3). Data were analyzed by unpaired two-tailed Student's *t*-test; *, $P<0.05$; **, $P<0.01$; ***, $P<0.001$; *ns*, $p>0.05$.

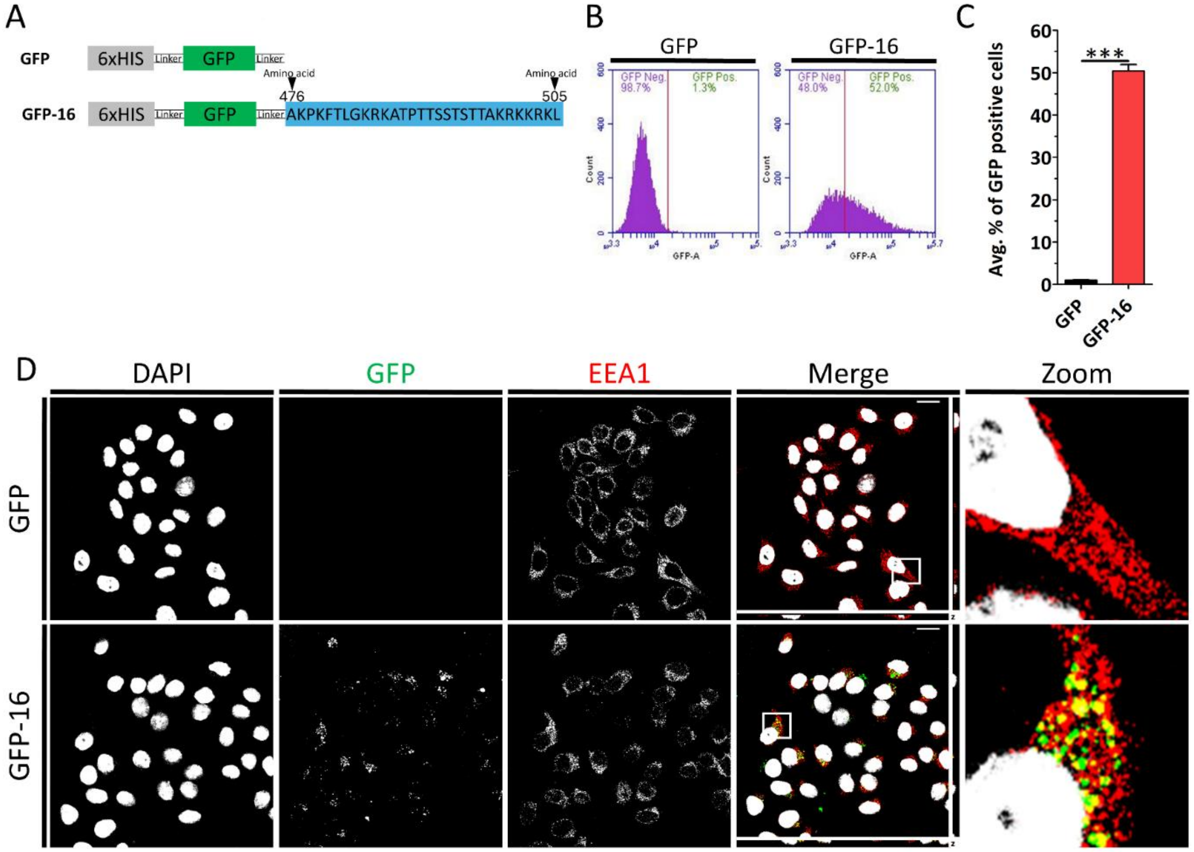


**FIGURE S2 |** GFP-16 enters HeLa cells. (A) Schematic representation of GFP and GFP-16 fusion protein construct. (B) HeLa cells were treated with 3 μM of GFP or GFP-16 at 37 ºC for 2 h. Internalization was analyzed using flow cytometry. Representative histograms show the percentage of GFP-positive cells. (C) Quantification of average percentage of GFP-positive cells. (D) Confocal fluorescence microscopy images of HeLa cells treated with 3 μM GFP or GFP-16 for 2 h at 37 ºC. Gray corresponds to DAPI (nuclear stain), green corresponds to GFP, and red corresponds to EEA1 (early endosomal marker). Scale bar: 20 μm. Data are represented as mean ± SEM (n=3). Data were analyzed by unpaired two-tailed Student's *t*-test; *, *P*<0.05; **, *P*<0.01; ***, *P*<0.001; *ns*, p>0.05.

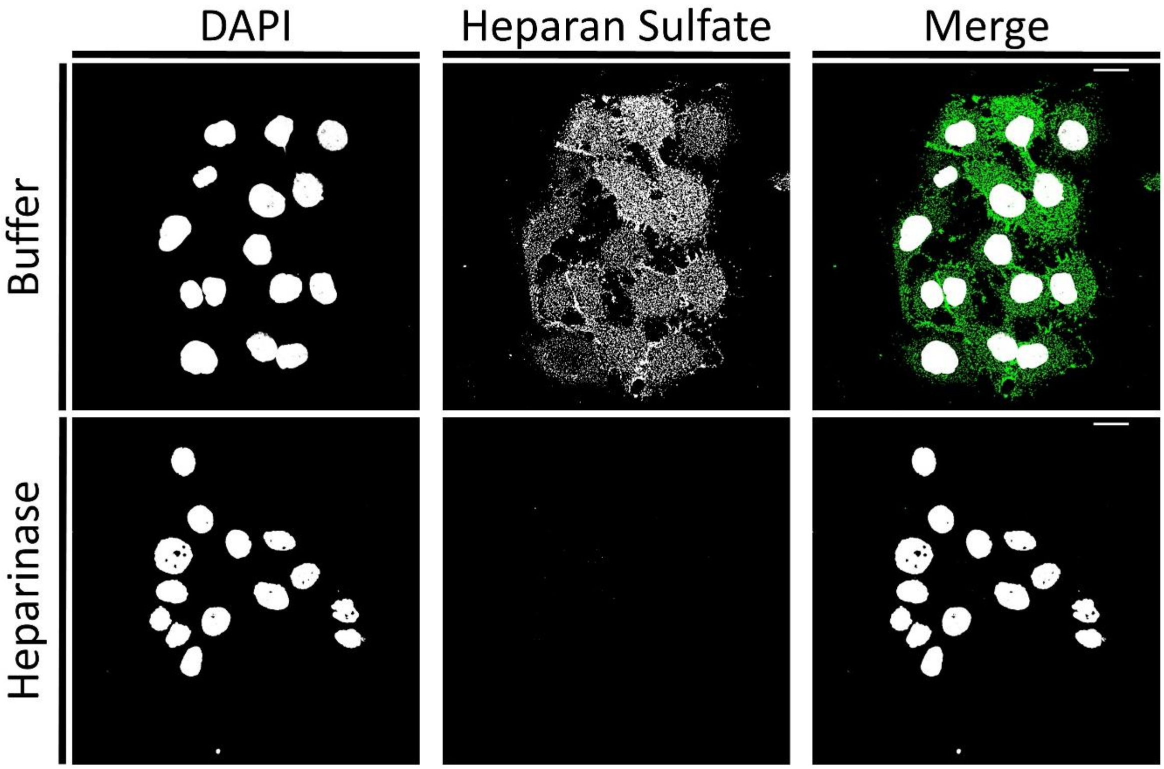


**FIGURE S3 |** Heparinase removes cell surface heparan sulfate. HaCaT cells treated with Buffer (control) or Heparinase for 2 h. at 37 ºC. After the treatment, the cells were washed and incubated in fresh media for an additional 2 h at 37 ºC. The cells were then fixed, immunostained for heparan sulfate, and imaged with confocal fluorescence microscopy. Gray corresponds to DAPI (nuclear stain) and green corresponds to heparan sulfate.

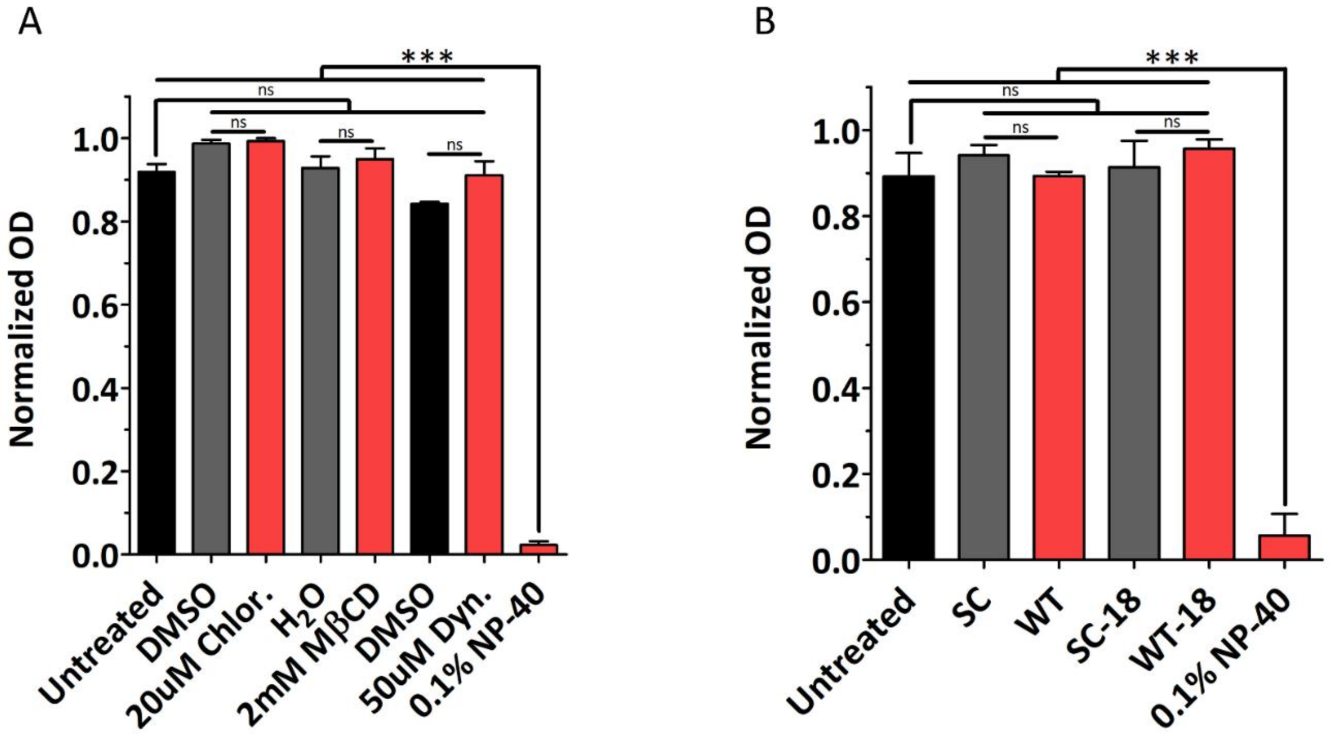


**FIGURE S4 |** Cytotoxicity assay of cells treated with endocytic inhibitors and peptides. (A) HaCaT cells were treated with the given concentration of endocytic inhibitor for 4 h at 37 ºC. (B) HaCaT cells were treated with 18 μM of each peptide for 48 h at 37 ºC. Data are represented as mean ± SEM (n=3). Data were analyzed by one-way ANOVA (Bonferroni's multiple comparisons test; *, P<0.05; **, P<0.01; ***, P<0.001; ns, p>0.05).

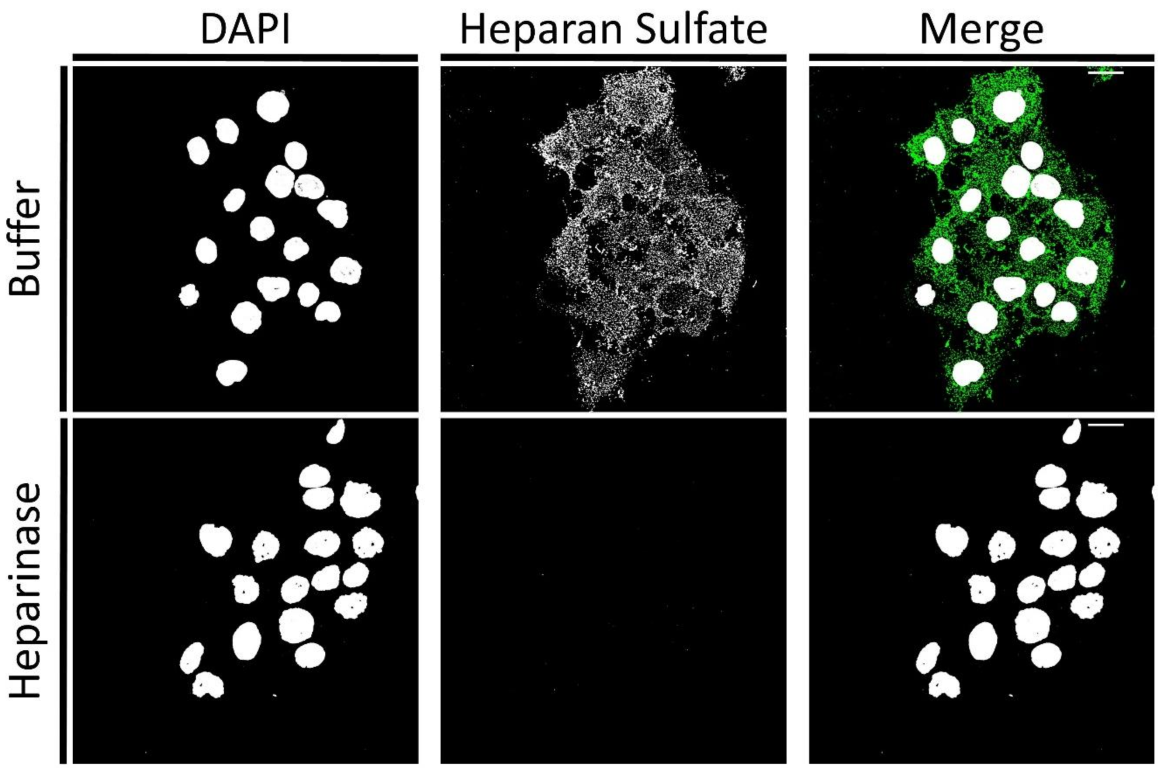


**FIGURE S5** | Heparinase removes cell surface heparan sulfate. HaCaT cells treated with Buffer (control) or Heparinase for 2 h at 37 ºC. After the treatment, the cells were washed and incubated in fresh media for an additional 2 h at 37 ºC. The cells were then fixed, immunostained for heparan sulfate, and imaged with confocal fluorescence microscopy. Gray corresponds to DAPI (nuclear stain) and green corresponds to heparan sulfate.
